# Longitudinal per-lesion *in vivo* imaging reveals allele-dependent resistance evolution in EGFR-mutant lung cancer

**DOI:** 10.64898/2026.04.17.717686

**Authors:** Eva Cabrera San Millan, Daniele Panetta, Paolo Armanetti, Mauro Quaglierini, Alessandro Zega, Raffaella Mercatelli, Emilia Bramanti, Luca Menichetti, Giorgia Maroni, Elena Levantini

**Author notes:** correspondance: Elena Levantini, PhD, Giorgia Maroni, PhD. co-last (Giorgia Maroni, Elena Levantini).

## Abstract

Acquired resistance to targeted therapies remains an inevitable outcome in EGFR-mutant non-small cell lung cancer, yet the spatiotemporal dynamics through which resistant clones emerge and evolve *in vivo* remain incompletely understood. In particular, how distinct oncogenic EGFR alleles shape evolutionary trajectories under therapeutic pressure within native tumor microenvironments remains unclear.

Here, we establish a longitudinal *in vivo* imaging framework to resolve tumor evolution at single-lesion resolution in genetically engineered mouse models (GEMMs) harboring three clinically relevant EGFR mutations: exon 19 deletion (EGFR^D19^), L858R (EGFR^LR^), and L858R/T790M (EGFR^LT^). Using high-resolution micro-computed tomography, three-dimensional reconstruction, and per-lesion volumetric tracking, we quantitatively map tumor growth dynamics, therapeutic response, and resistance emergence over time in individual lesions within the same animal.

We find that EGFR alleles impose distinct evolutionary trajectories. EGFR^LT^ -driven tumors exhibit shorter latency and early emergence of lesions with intrinsic resistance to osimertinib. In contrast, EGFR^D19^ and EGFR^LR^ tumors show slower growth kinetics, more homogeneous initial responses, and delayed acquisition of resistance during prolonged treatment.

Importantly, longitudinal per-lesion imaging reveals marked spatial heterogeneity across all genotypes. Within the same lung microenvironment, individual lesions undergo complete regression, sustained response, or progressive growth, reflecting parallel and spatially distinct evolutionary trajectories. These divergent behaviors emerge despite shared systemic therapy and identical host environment, underscoring lesion-intrinsic and genotype-dependent constraints on evolution.

Together, these findings identify oncogenic EGFR genotype as a key determinant of the temporal and spatial architecture of resistance evolution under targeted therapy. More broadly, we provide a quantitative framework to resolve tumor evolution *in vivo* at lesion-level resolution, applicable to dissecting spatiotemporal dynamics of tumor growth and therapeutic response across oncogene-driven cancers.

## INTRODUCTION

Non-small cell lung cancer (NSCLC) accounts for ∼85% of lung cancer cases and remains the leading cause of cancer-related mortality (*1, 2*). Lung adenocarcinoma (LUAD), the most common subtype, is frequently driven by activating mutations in the epidermal growth factor receptor (EGFR), occurring in ∼15-20% of Caucasian patients and associated with poor prognosis and reduced therapeutic efficacy (*3-5*).

The most prevalent EGFR alterations are in-frame deletions in exon 19 (Del19) and the L858R missense mutation in exon 21, which together account for ∼85% of common EGFR activating mutations (*4, 6, 7*). Patients harboring these mutations are initially sensitive to tyrosine kinase inhibitors (TKIs), which represent the standard first-line therapy (*4, 7*). Among them, osimertinib, a third-generation TKI, significantly improves progression-free survival (PFS) and overall survival (OS) compared with earlier-generation inhibitors (*7, 8*), as it targets both activating EGFR mutations and T790M-mediated resistance that emerges during prior TKI therapy.

Despite initial responses, acquired resistance to osimertinib is inevitable and emerges through highly heterogeneous mechanisms. Clinical evidence indicates that resistance patterns differ across EGFR genotypes, with variation in progression kinetics, secondary alterations, and clinical outcomes, suggesting an allele-dependent component of therapeutic escape (*9-14*). Consistently, clinical and genomic analyses have demonstrated that osimertinib resistance is mechanistically diverse, with heterogeneous resistance mechanisms and clinical response patterns across patients, including on-target EGFR alterations and off-target bypass signaling events across patients and tumor sites (*4, 15-19*). Importantly, resistance arises through selection of pre-existing and/or therapy-emergent resistant clones under therapeutic pressure, as demonstrated in preclinical and patient-derived models of osimertinib resistance (*20*).

In parallel, converging preclinical and clinical evidence supports the existence of a drug-tolerant persister state in EGFR-mutant NSCLC, in which a fraction of tumor cells survive transient EGFR inhibition through non-genetic, reversible adaptive programs (*20-22*), thereby providing a reservoir for subsequent clonal expansion and disease relapse.

Spatial intratumor heterogeneity further contributes to variable tumor composition, reflecting divergent clonal architecture and branched evolutionary trajectories, as demonstrated by multiregion sequencing studies in solid tumors (*23*). Such spatial heterogeneity is further amplified by branched evolutionary processes, leading to regionally distinct subclones with differential drug sensitivity within the same lesion (*23-25*). Consistently, heterogeneous clinical patterns of mixed response and progression have been reported in EGFR-targeted NSCLC, reflecting lesion-specific evolutionary trajectories (*26-29*).

However, the determinants of mutation-associated resistance remain incompletely understood, and the spatiotemporal dynamics of tumor response and clonal evolution during treatment are not fully characterized, limiting our ability to anticipate and intercept resistance progression (*4, 30-34*).

Importantly, therapeutic response is not uniform across EGFR genotypes. Exon 19 deletions are generally associated with higher objective response rates (ORR) and longer progression-free survival (PFS) compared with L858R mutations, both in first-line and broader clinical cohorts (*35, 36*) reflecting biological and clinical heterogeneity among EGFR mutation subtypes (*37*). Subgroup analyses further indicate that specific Del19 variants, particularly those initiating at E746, are associated with improved outcomes compared with other deletion patterns (*38*), highlighting clinically relevant heterogeneity within EGFR exon 19 deletions. In T790M-positive cohorts, osimertinib also demonstrates improved efficacy in Del19 compared with L858R tumors (*9*).

To dissect these allele-specific effects, we employed genetically engineered mouse models (GEMMs) carrying clinically relevant EGFR mutations: exon 19 deletions (EGFR^D19^), L858R (EGFR^LR^), and L858R/T790M (EGFR^LT^). Mice were treated with osimertinib until resistance developed, generating osimertinib-resistant (OsiR) tumors. Tumor progression and therapeutic response were monitored longitudinally using high-resolution micro-computed tomography (micro-CT), allowing three-dimensional (3D) reconstruction, precise quantification of tumor burden, and lesion-specific tracking over time.

This longitudinal *in vivo* platform enables allele-specific comparison of tumor growth kinetics, drug response, and resistance evolution under clinically relevant conditions of EGFR-targeted therapy. Unlike conventional xenograft or *in vitro* systems, it preserves spatiotemporal tumor architecture and multiclonal heterogeneity, enabling mechanistic dissection of resistance emergence in EGFR-mutant NSCLC.

Overall, this framework enables the investigation of how EGFR genotype, clonal architecture, and spatially distinct tumor subpopulations jointly shape the timing, localization, and evolutionary routes of resistance under sustained osimertinib exposure.

## RESULTS

### Generation and Longitudinal Monitoring of Osimertinib-Resistant EGFR GEMMs Reveals Allele-Specific Differences in Tumor Growth and Resistance Development

To investigate the dynamics of osimertinib resistance *in vivo*, we generated OsiR GEMMs across three distinct EGFR mutant backgrounds (EGFR^D19^, EGFR^LR^, and EGFR^LT^) (**Fig. 1**), inducing lung tumors via doxycycline-dependent activation of mutant EGFR and monitoring tumor growth longitudinally by high-resolution micro-CT.

**Figure 1.**
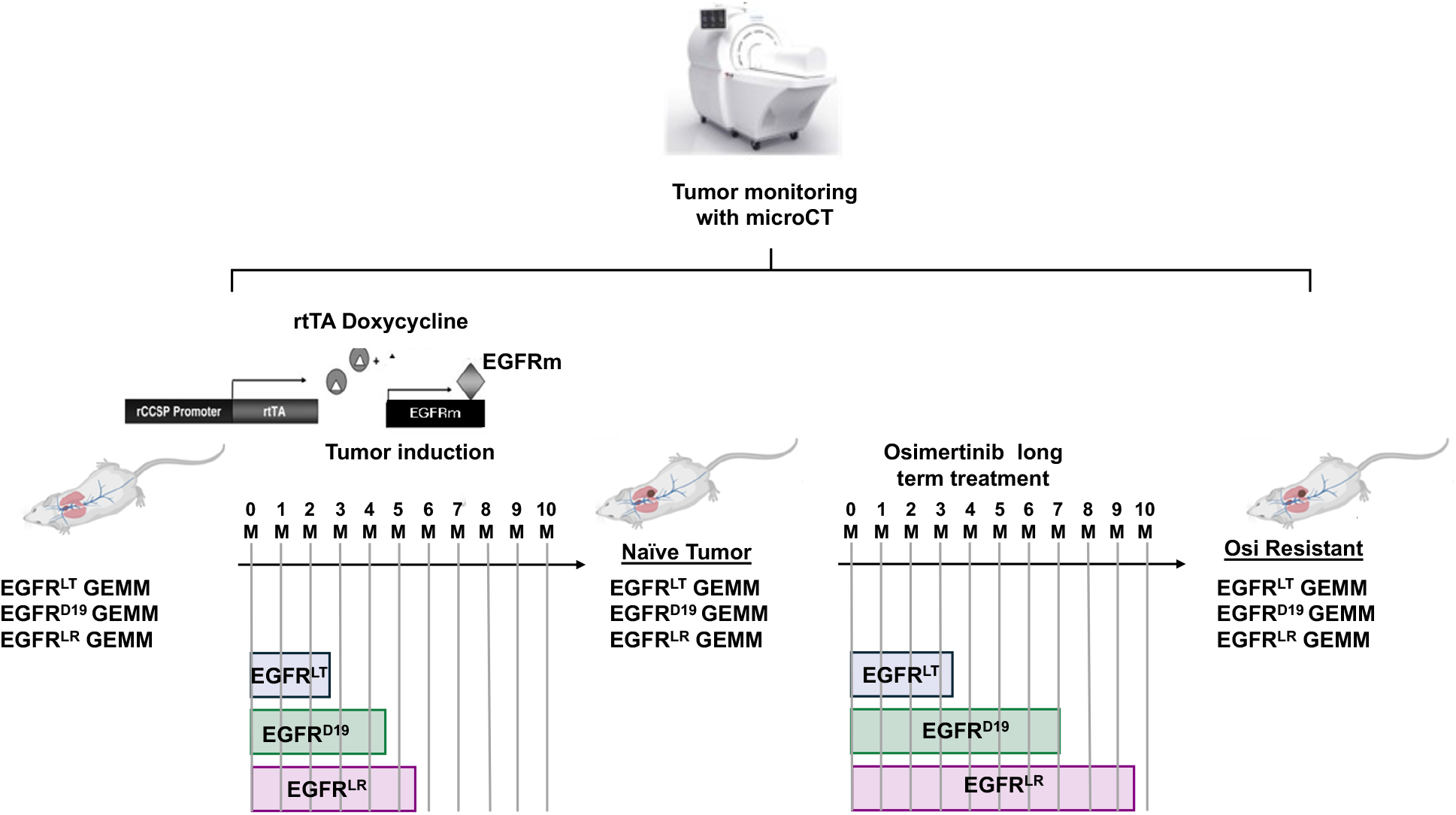
Experimental strategy for the generation of osimertinib-resistant EGFR GEMMs. Schematic overview of tumor induction and treatment timeline in EGFR-driven GEMMs. Tumors were induced by doxycycline-dependent expression of mutant EGFR alleles, followed by longitudinal monitoring with high-resolution micro-CT. M indicates months. After baseline imaging of naïve tumors (week 0), mice undergo continuous osimertinib treatment, leading to the emergence of osimertinib-resistant tumors. The timeline illustrates the duration of treatment and the micro-CT imaging schedule for the EGFR^LT^, EGFR^D19^, and EGFR^LR^ models.

Tumor initiation exhibited marked allele-specific differences: EGFR^LT^ tumors became detectable within 2-3 months of induction, EGFR^D19^ tumors after 4-5 months, and EGFR^LR^ tumors showed the longest latency, reaching comparable volumes only after 5-6 months. These findings demonstrate that the oncogenic EGFR allele strongly influences tumor initiation kinetics *in vivo*, establishing distinct evolutionary baselines prior to therapeutic pressure.

Once tumors reached measurable size, mice were treated with osimertinib to induce acquired resistance. The timing of resistance emergence, defined as tumor re-growth during continuous therapy, varied substantially among genotypes (**Fig. 1**). EGFR^LT^ tumors developed resistance after ∼3.5 months, EGFR^D19^ tumors after ∼7 months, and EGFR^LR^ tumors after ∼9.5 months of therapy. Notably, in this study, the genotype associated with the shortest tumor latency (EGFR^LT^) also displayed the fastest acquisition of resistance, whereas genotypes with delayed tumor formation exhibited prolonged sensitivity to osimertinib. Together, these data indicate that distinct oncogenic EGFR mutations drive allele-specific evolutionary trajectories under therapeutic pressure *in vivo*, shaping both tumor growth kinetics and resistance timing.

### Longitudinal Response Dynamics in EGFR^D19^ and EGFR^LR^ Tumors

Longitudinal micro-CT imaging was used to monitor tumor dynamics during osimertinib treatment. Tumor volumes were calculated relative to the pre-treatment (naïve) state, and 3D reconstructions were generated at defined stages to visualize lesion distribution and dynamics.

In both EGFR^D19^ and EGFR^LR^ models, 3D representative reconstructions revealed comparable patterns of initial tumor response followed by resistance development (**Figs. 2a, 3a**). At baseline (naïve stage), multiple discrete lesions were detectable throughout the lungs.

**Figure 2.**
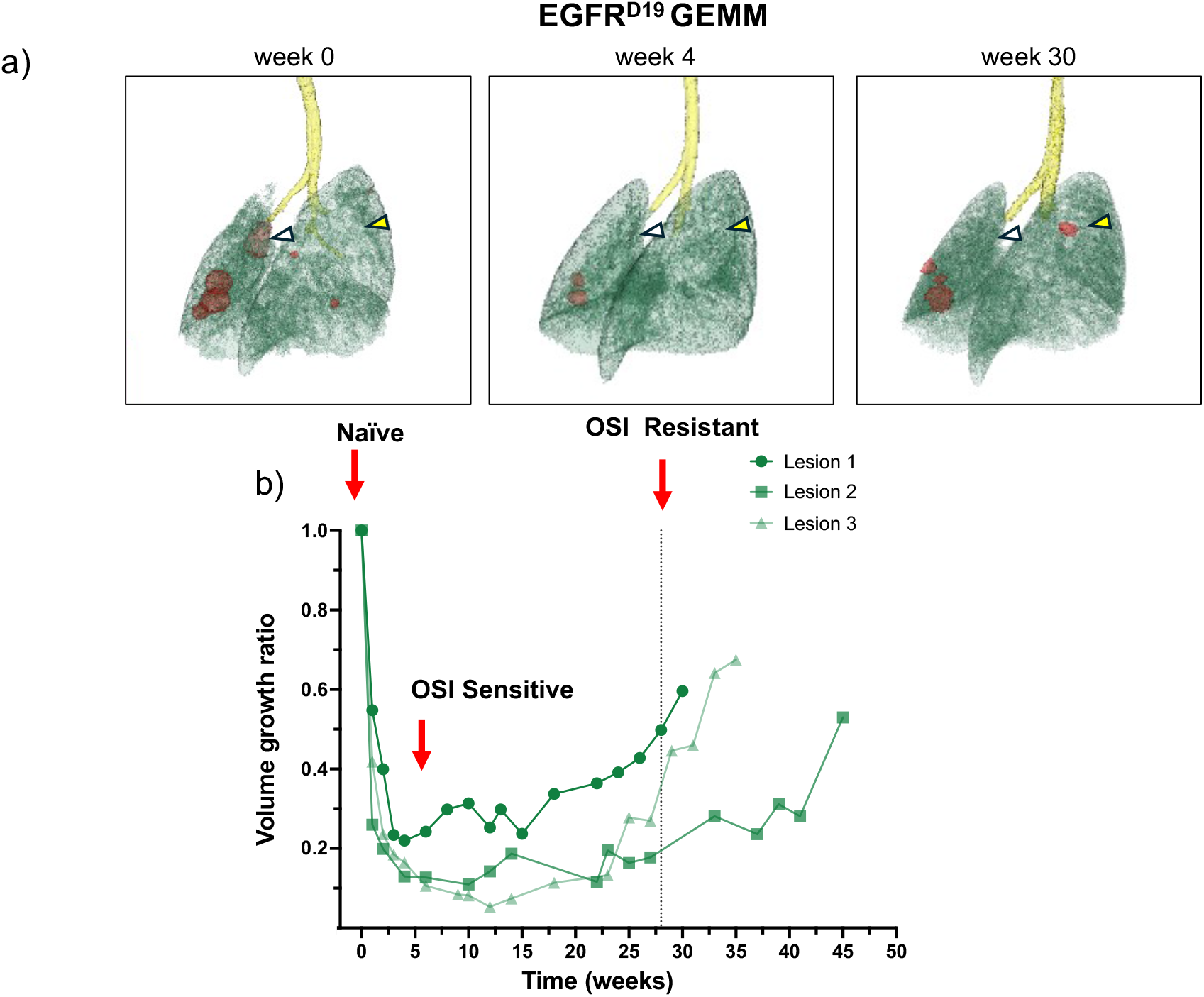
Generation of osimertinib resistant EGFR^D19^ GEMMs. **(a)** Representative micro-CT-based 3D reconstructions showing lung tissue (green), tumor lesions (red), and trachea (yellow) at baseline (week 0, naïve stage), during response to osimertinib (week 4), and after acquisition of resistance at the end of treatment (week 30). White arrows indicate lesions that regress completely and do not re-grow, whereas yellow arrows mark lesions that emerge during the resistant phase. **(b)** Tumor volume growth kinetics of individual lesions showing initial regression followed by tumor re-growth upon acquired resistance. Each curve indicates one representative lesion per mouse (*n* = 3) that remained measurable throughout treatment. Red arrows point to naïve, responding (OsiS), and resistant (OsiR) lesions.

**Figure 3.**
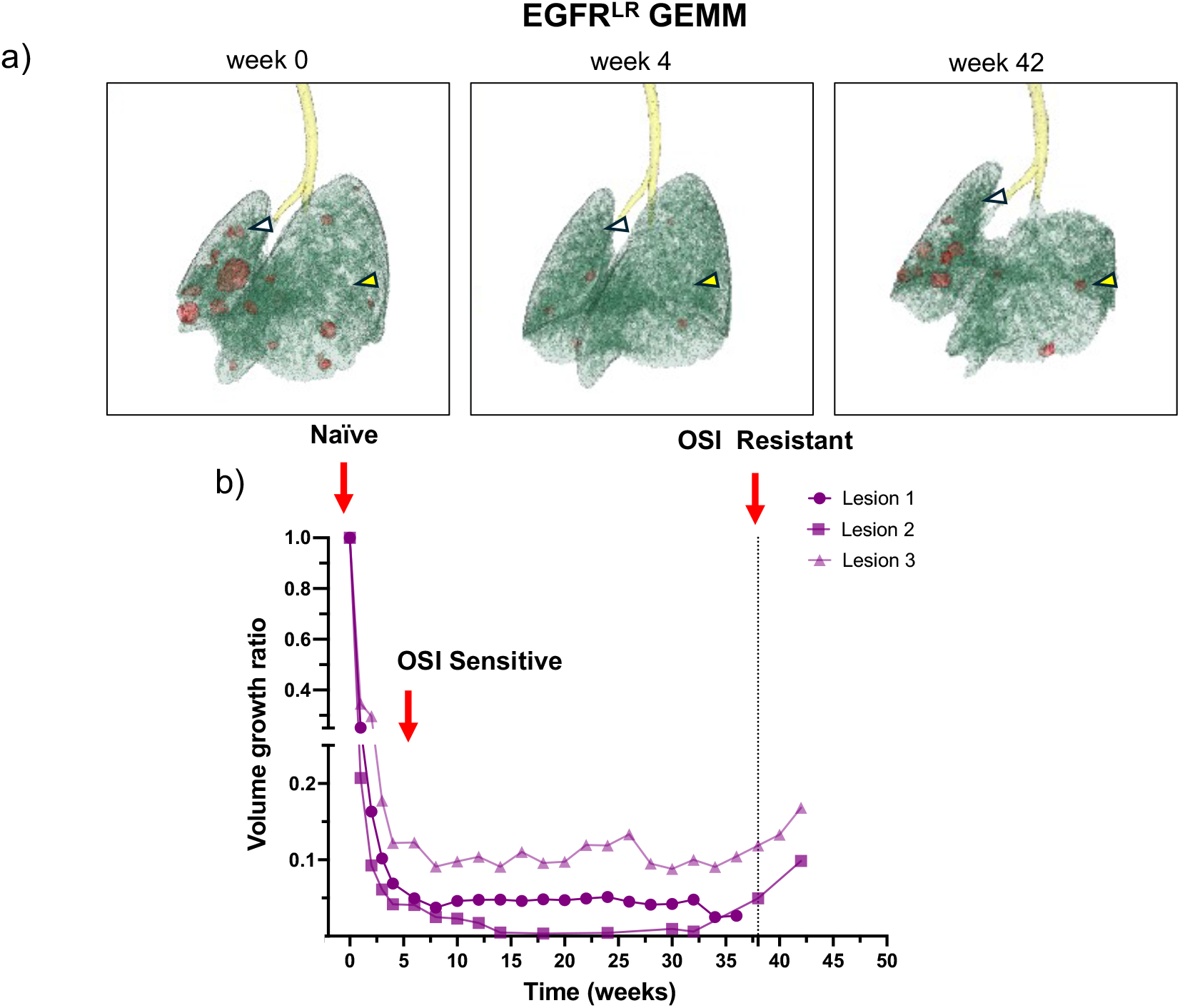
Generation of osimertinib-resistant EGFR^LR^ GEMMs. **(a)** Representative micro-CT-based 3D reconstructions showing lung tissue (green), tumor lesions (red), and trachea (yellow) at baseline (week 0, naïve stage), during response to osimertinib (week 4), and after acquisition of resistance at the end of treatment (42 weeks). White arrows indicate lesions that regress completely and do not re-grow, whereas yellow arrows mark lesions that emerge during the resistant phase. **(b)** Tumor volume growth kinetics of individual lesions showing initial regression followed by tumor re-growth upon acquired resistance. Each curve indicates one representative lesion per mouse (n = 3) that remained measurable throughout treatment. Red arrows point to naïve, responding (OsiS), and resistant (OsiR) lesions.

Following treatment initiation, most lesions exhibited rapid volumetric regression within the first four weeks. A subset of lesions regressed completely and remained undetectable throughout prolonged therapy **(Figs. 2a, 3a**), suggesting durable elimination of specific tumor clones. Conversely, additional lesions became detectable at later time points **(Figs. 2a, 3a**), indicating the emergence or expansion of previously undetectable tumor populations during therapy.

For quantitative assessment, one representative lesion per mouse (n=3) that remained measurable throughout treatment was longitudinally tracked. In the EGFR^D19^ GEMMs, lesions underwent ∼75-80% volume reduction by week 4, followed by stabilization until ∼week 25-28, when tumor re-growth marked the onset of acquired resistance (**Fig. 2b**).

Although resistance emerged around weeks 25-28, treatment was maintained beyond this time point to allow further expansion of resistant lesions, ensuring sufficient tumor volume at sacrifice for downstream analyses.

In EGFR^LR^ GEMMs, initial regression was ∼80-90% by week 4, with tumor volumes remaining largely stable over an extended period (weeks 4-38). During this phase, some lesions decreased slightly or remained largely stable, while others became nearly undetectable, reflecting sustained tumor control under continuous therapy. Measurable re-growth began after week 38, and treatment continued until week 42 to allow resistant tumors to reach sufficient volume for downstream analyses (**Fig. 3b**).

For one of the three EGFR^LR^ mice, quantification was interrupted toward the end of treatment because the initially tracked lesion was no longer clearly detectable. Nonetheless, new dense regions and signs of localized inflammation emerging between weeks 38-42, particularly in the left lung lobe, likely corresponded to new resistant lesions (**Supplementary Fig. 1**), confirming the development of resistance.

These kinetics were consistent across animals, underscoring the reproducibility of treatment response and resistance timing within each model.

Together, EGFR^D19^ and EGFR^LR^ tumors displayed broadly similar response trajectories, characterized by robust initial sensitivity to osimertinib and durable regression, although resistance emerged later in EGFR^LR^ tumors, indicating subtle allele-specific differences in temporal dynamics.

Overall, combined 3D reconstructions and longitudinal volumetric tracking provide a robust platform for quantitative lesion-level assessment of therapeutic response and resistance evolution *in vivo*.

**Supplementary Fig. 1.**
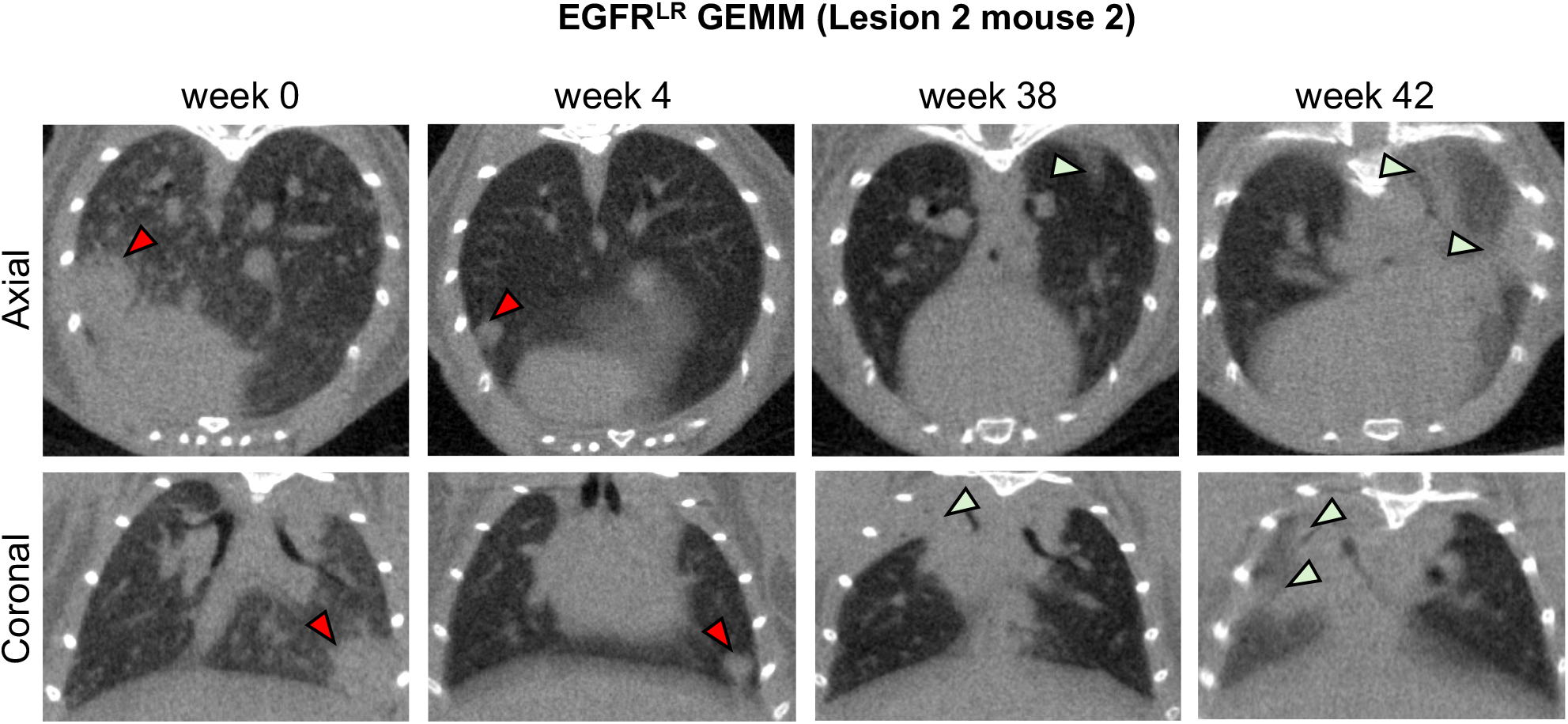
Representative 2D micro-CT images of the OsiR EGFR^LR^ model. Representative 2D micro-CT images in coronal (top) and axial (bottom) views of one of n=3 mice from the OsiR EGFR^LR^ GEMM group. Images are shown at baseline (week 0, naïve stage), during response to osimertinib (week 4), and after acquisition of resistance at the end of treatment (weeks 38 and 42). Red arrows indicate lesions that were quantified, which initially responded to treatment but were later lost. Green arrows mark areas of inflammation and potential emerging resistant lesions during the resistant phase, which are partially obscured by inflammation and therefore difficult to quantify. This figure illustrates the temporal dynamics of lesion regression, the emergence of resistant clones, and challenges in quantifying lesions during inflammation, complementing 3D reconstructions shown in Figure 3.

### Intrinsic and heterogeneous resistance trajectories in EGFR^LT^ tumors

By contrast, the EGFR^LT^ GEMMs revealed markedly heterogeneous patterns of response and resistance (**Fig. 4a**). At baseline (week 0), multiple discrete lesions were detected throughout the lungs. Following four weeks of osimertinib treatment, some lesions exhibited rapid volumetric regression (∼70-80%) (**Fig. 4b**), but early re-growth began between weeks 10-15 (**Figs. 4b, 4c**), substantially earlier than in EGFR^D19^ and EGFR^LR^ models.

**Figure 4.**
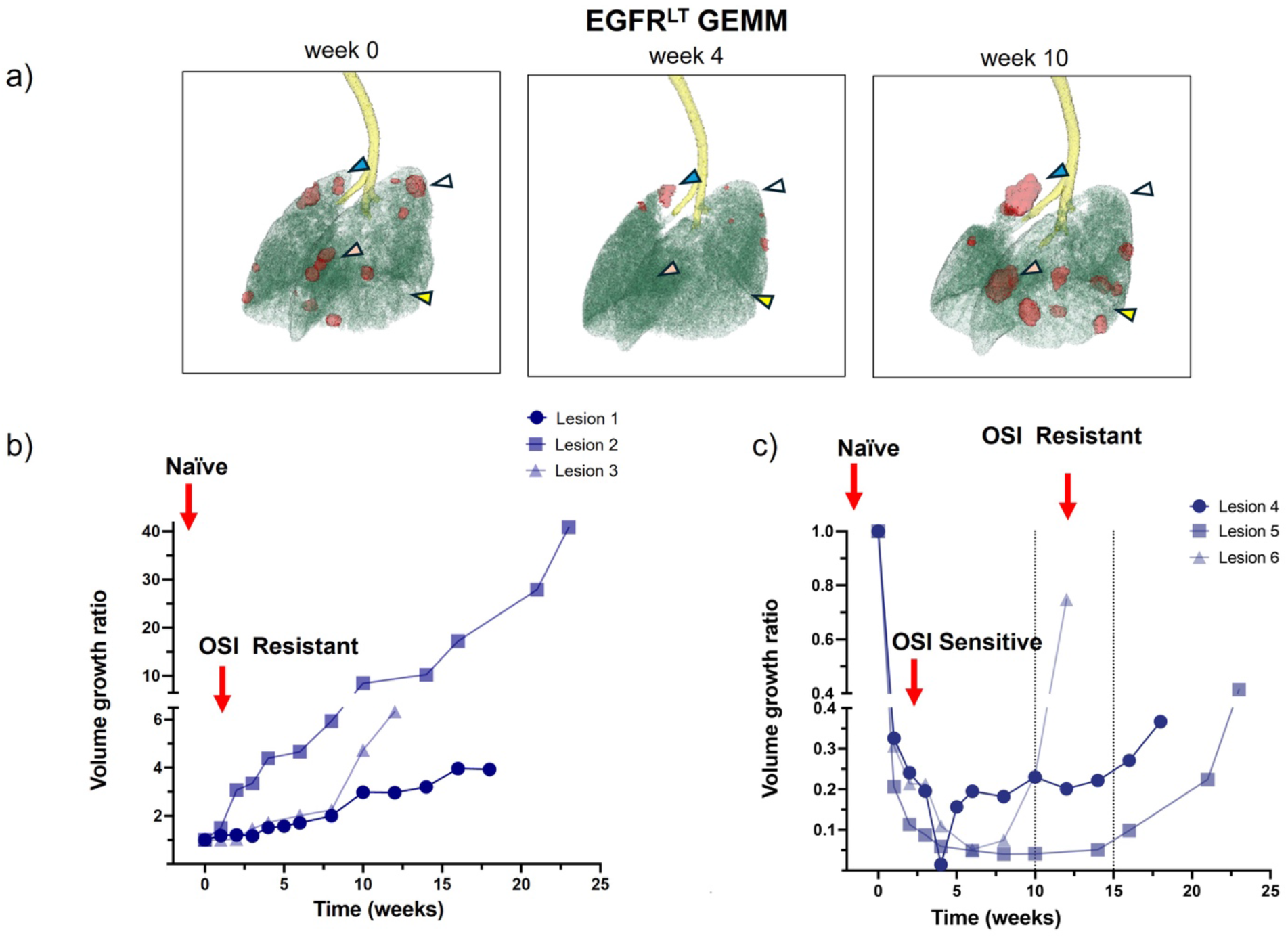
Generation of osimertinib-resistant EGFR^LT^ GEMMs. **(a)** Representative micro-CT-based 3D reconstructions showing lung tissue (green), tumor lesions (red), and trachea (yellow) at baseline (week 0, naïve stage), during response to osimertinib (week 4), and after acquisition of resistance (week 10). White arrows indicate lesions that respond to treatment and do not re-grow, whereas yellow arrows represent lesions that emerge during the resistant phase. Blue arrows indicate intrinsically resistant lesions. **(b-c)** Tumor volume growth kinetics of individual lesions showing either non-responsive growth consistent with intrinsic resistance **(b)** or initial regression followed by tumor re-growth upon acquired resistance **(c)**. Each curve indicates one representative lesion per mouse (n = 3). Red arrows point to naïve, responding (OsiS), and resistant (OsiR) lesions.

A subset of lesions regressed completely and remained undetectable during prolonged therapy, suggesting durable eradication of specific tumor clones. Simultaneously, other lesions demonstrated intrinsic resistance, showing no measurable regression and progressing continuously despite therapy (blue arrows, **Fig. 4a**). New lesions emerged only at later time points, highlighting pronounced spatial and clonal heterogeneity within individual lungs.

For quantitative assessment (**Fig. 4b-c**), at least two representative lesions per mouse (n=3) that remained measurable throughout treatment were longitudinally tracked. This analysis captured two distinct resistance trajectories: intrinsically resistant lesions exhibiting continuous growth during therapy (**Fig. 4b**), and initially responsive lesions showing early regression followed by subsequent re-growth (**Fig. 4c**).

Collectively, EGFR^LT^ GEMMs represent a highly heterogeneous model of osimertinib response, characterized by the coexistence of sensitive and intrinsically resistant clones and complex spatial dynamics within the lungs.

These observations emphasize the importance of lesion-level monitoring for understanding the mechanisms underlying therapeutic escape.

## DISCUSSION

Understanding how resistance to EGFR-targeted therapies emerges remains a central challenge in EGFR-mutant lung cancer. Third-generation EGFR inhibitors such as osimertinib have substantially improved outcomes in EGFR-mutant NSCLC, yet resistance inevitably develops. Therapeutic escape is driven by heterogeneous and often co-occurring mechanisms reflecting complex temporal, spatial, and clonal evolutionary dynamics within tumors (*15, 16, 39*).

In this study, we established a longitudinal *in vivo* platform to investigate osimertinib resistance using GEMMs harboring clinically relevant EGFR mutations (EGFR^D19^, EGFR^LR^, and EGFR^LT^). By integrating high-resolution micro-CT imaging with lesion-specific volumetric tracking, we longitudinally monitored tumor initiation, therapeutic response, and resistance at both whole-lung and lesion resolution. This approach enabled direct reconstruction of mutation-specific evolutionary trajectories under sustained therapeutic pressure.

Our data demonstrate that EGFR mutational context strongly influences tumor growth kinetics and therapeutic response. EGFR^LT^ tumors exhibited the most rapid onset and the earliest resistance emergence, whereas EGFR^D19^ and EGFR^LR^ tumors showed slower-growing tumors and prolonged therapeutic control. These findings indicate that EGFR alleles impose distinct evolutionary constraints on tumor progression, consistent with previously reported mutation-dependent clinical outcomes and differential responses to EGFR-targeted therapies in patients (*8-10, 36, 39*).

Lesion-level analyses revealed marked intratumoral heterogeneity, particularly in EGFR^LT^ tumors, where intrinsically resistant lesions coexisted with initially responsive lesions that later relapsed. In contrast, EGFR^D19^ and EGFR^LR^ tumors displayed more homogeneous response patterns characterized by robust initial regression and delayed re-growth. These dynamics support a model in which resistance emerges through selection of pre-existing resistant subclones and/or therapy-driven evolutionary adaptation under drug pressure (*40-44*).

From an evolutionary perspective, our results suggest a trade-off between tumor growth rate and duration of response. Rapidly proliferating tumors may be associated with increased opportunities for clonal diversification, accelerating therapeutic escape, whereas slower-growing tumors maintain more constrained evolutionary trajectories and prolonged drug sensitivity.

Importantly, conventional clinical readouts such as bulk imaging or circulating tumor DNA may obscure lesion-specific dynamics. Our longitudinal imaging and per-lesion analysis framework enables resolution of spatially distinct evolutionary trajectories that are not accessible in routine clinical monitoring.

Collectively, our preclinical models demonstrate that EGFR genotype in the therapy-naïve setting strongly influences tumor behavior and resistance evolution. Longitudinal lesion-level tracking revealed that resistant clones emerge in spatially distinct regions of the lung, underscoring the spatial and temporal complexity of tumor evolution during treatment. In the EGFR^LT^ model, intrinsically resistant lesions coexisted with initially sensitive lesions that later relapsed, illustrating how heterogeneous tumor subclones can follow distinct evolutionary trajectories within the same organism. These observations support a model in which both pre-existing resistant populations and therapy-induced selective pressures contribute to disease progression.

From a translational perspective, this platform is conceptually aligned with the emerging notion of functional avatar models in oncology. By enabling early, spatially resolved tracking of tumor evolution under therapeutic pressure, GEMMs combined with longitudinal imaging may serve as dynamic preclinical avatars to investigate disease evolution under clinically relevant selective pressures and to inform adaptive therapeutic strategies. Importantly, GEMMs preserve an intact and immunocompetent tumor microenvironment, thereby maintaining continuous tumor-immune-stroma interactions during evolution. In line with our previous single-cell analyses in EGFR-driven lung cancer models (*7*), these data support a model in which resistance emerges within a physiologically relevant ecosystem rather than in tumor-intrinsic isolation.

By highlighting mutation-specific growth kinetics, pre-existing clonal diversity, and lesion-level heterogeneity, this study underscores the importance of modeling diverse EGFR mutational contexts. Our allele-specific GEMMs combined with longitudinal imaging provide a powerful experimental platform to study resistance evolution and to evaluate therapeutic strategies aimed at delaying or preventing resistance.

Finally, the modular nature of this imaging-based framework is readily adaptable to additional therapeutic contexts, including combination regimens and sequential treatment strategies, enabling systemic investigation of how different therapeutic pressures shape evolutionary trajectories.

In summary, our study reveals that EGFR allele context dictates the temporal and spatial dynamics of resistance evolution under osimertinib therapy, while intratumoral lesion-level heterogeneity drives the coexistence of spatially segregated and evolutionarily distinct (sensitive and intrinsically resistant lesions) tumor subpopulations within the same tumor ecosystem.

## MATERIAL AND METHODS

### Establishment of GEMMs colonies and generation of OsiR GEMMs models

Bitransgenic mice were generated by crossing TetO-regulated human EGFR mutant lines (EGFR^LT^, EGFR^D19^, and EGFR^LR^) with mice expressing reverse tetracycline transactivator under the Clara Cell Secretory Protein promoter (CCSP-rtTA)(*45-47*). In this system, mutant EGFR expression is restricted to pulmonary epithelial cells and induced upon doxycycline administration, as previously described (*7, 48-50*).

To initiate transgene expression and tumorigenesis, 8-10-week-old mice were fed a doxycycline-impregnated diet continuously. Lung tumor development was monitored by high-resolution micro-CT imaging performed every two weeks, and baseline tumor burden was established prior to treatment initiation.

Once tumors were detectable by imaging, mice were treated with osimertinib (10 mg/kg) administered by oral gavage twice weekly until acquired resistance was induced. A minimum of three mice per EGFR genotype were included for the generation of resistant models. Treatment continued until evidence of tumor re-growth, consistent with the emergence of osimertinib resistance, was observed. Tumor burden was quantified by serial micro-CT imaging, initially weekly and then every two weeks, enabling lesion-specific longitudinal tracking.

All experiments were performed in compliance with ethical regulations and approved by the Italian Ministry of Health (488/2022-PR).

### High resolution micro-CT Imaging

All micro-CT scans were performed with the IRIS scanner (Inviscan, Strasbourg, France). Prior to imaging, mice were pre-anaesthetized with isoflurane in an induction chamber and subsequently positioned on the imaging bed, where anesthesia was maintained using a continuous mixture of isoflurane and oxygen delivered via a nose mask.

Scanning parameters were as follows: 65 kVp, 1 mA, 1280 projections over 360°, 70 ms of exposure time per projection, and a total scan time of 90 seconds. Images were corrected for beam hardening artifacts and reconstructed using cone-beam filtered backprojection (FBP) onto a 3D matrix of 600×600×1458 isotropic voxels, with a voxel size of 77 μm. Reconstructed datasets were exported in DICOM format for archiving and downstream quantitative image analysis. Given the stable depth of sedation and regular free breathing pattern, respiratory gating was not required to obtain high-quality reconstructions of pulmonary lesions. In addition, the strong intrinsic contrast between solid tumor lesions and surrounding lung parenchyma allowed imaging without the use of exogenous contrast agents.

### Tumor lesion quantification

Tumor volumes were quantified from DICOM micro-CT images using 3D Slicer (v5.8.0). For each mouse, one or two representative lesions were selected and longitudinally tracked across all time points. A dedicated segment was created for each lesion. Segmentation was performed using the Sphere Brush tool under masking conditions, applying the “Editable intensity range” option to restrict editing to predefined lesion intensity thresholds. Lesions were manually refined on a slice-by-slice basis to ensure accurate volumetric delineation and were systematically reviewed in both coronal and axial planes to confirm correct anatomical localization. Volumetric measurements were extracted using the Segment Statistics module, with identical segmentation thresholds applied consistently across all time points. The primary endpoint was labelmap (LM) volume (mm^3^). Lesion volumes were recorded at each time point until the development of resistance. Tumor growth volume dynamics were calculated relative to baseline (week 0, naïve tumor) and plotted for each EGFR mutant group.

### Three-dimensional lung reconstruction

Three-dimensional reconstruction and segmentation were performed using 3D Slicer (v5.8.0). For each EGFR-mutant GEMM group (EGFR^LT^, EGFR^D19^, and EGFR^LR^), one representative mouse was selected, and representative treatment time points were analyzed: pre-treatment (week 0), during response to osimertinib (osimertinib-sensitive phase, week 4), and at the final time point corresponding to acquired resistance (osimertinib-resistant phase), also corresponding to end of treatment.

DICOM micro-CT images were analyzed using the Segment Editor module. For lung and trachea segmentation, a new segment was created, and the Threshold tool was applied using an intensity range of - 1024 to -300 Hounsfield Units (HU). Selected regions were verified in axial and coronal planes to ensure accurate inclusion of both lung lobes and trachea. Segmentation refinement was performed using the “Grow from Seeds” tool to optimize boundary delineation and exclude surrounding soft tissues.

For tumor lesion reconstruction, a separate segment was generated. Thresholding was applied using an intensity range of -200 to +3900 HU to identify tumor regions. Manual refinement was performed in axial, coronal, and sagittal planes when necessary to ensure accurate lesion contouring. Closed surface representation was used to generate 3D volumetric reconstructions. For visualization, lung opacity was set to 0.6, lesion opacity to 1.0, and background structures to 0.25. All segmentation parameters were kept constant across samples and time points to ensure reproducibility and comparability.

### Statistics

Statistical analyses were performed using GraphPad Prism 10.4.1 (532). Specific details are provided in the figure legends.

## Acknowledgments

We sincerely thank Prof. Diego Cortinovis for the careful and thorough review of the manuscript, valuable and insightful suggestions.

The Euro-BioImaging research infrastructure located at the National Council of Research in Pisa has enabled open access to imaging technologies in the fields of biological and biomedical imaging, thereby providing support to this study.

**The authors declare that they have no competing interests**.

